# rpcFold: Residual Parallel Convolutional neural network to decipher RNA folding from RNA sequence

**DOI:** 10.1101/2024.08.26.609824

**Authors:** Nandita Sharma, Pralay Mitra

## Abstract

Precise secondary structure information offers deeper insights into the functionality of many RNA molecules. Earlier approaches, which primarily relied on the free-energy minimization, proved inadequate, as RNA often adopts complex folds, especially with increasing length. We present rpcFold, a residual parallel convolutional neural network-based model for RNA secondary structure prediction. From the nucleotide sequences, we compute a base-pairing possibility score using a locally weighted Gaussian function. It captures the intricate canonical and noncanonical pairing patterns, which enhances the modelling of short- and long-range dependencies. The features are mapped into an image that is the input to rpcFold. Our sliding window mechanism accommodates sequences of arbitrary length, and hence explicitly addresses the prediction of pseudoknots, which are often overlooked in prior works. The performance of rpcFold is demonstrated on nested and on non-nested (pseudoknot) base-pairs. While tested on within-family and cross-family benchmark datasets, rpcFold shows improved performance over existing state-of-the-art methods. Additionally, for long nucleotide sequences with complex pseudoknots, rpcFold achieves 71.1% F1-score in jointly predicting nested and non-nested pairs. Performance on unseen RNA families further confirms the robustness and adaptability of our approach. The prediction accuracy of rpcFold, particularly in long-range RNAs and pseudoknots, improves our understanding of RNA functions.

## 1. Introduction

RNA molecules play crucial roles in diverse cellular processes, such as genetic information transmission, transcriptional and post-transcriptional gene expression, and gene regulation [1]. Accurate structure information provides enhanced insights into the functionality of RNA. However, determining RNA structures is challenging because of their inherent flexibility and versatile structures [2]. Both secondary and tertiary structures of RNA hold the homology and evolutionary information, and contribute to functional analysis [3]. The tertiary structure of RNA is highly influenced by temperature and environmental factors, leading to poor stability [4]. Since RNA secondary structures are less complex and more stable than their corresponding tertiary structures, and contribute equally to functional predictions, researchers tend to focus more on RNA secondary structures. Also, the secondary structure is evolutionary more conserved than the primary sequence and is crucial in determining the function of many RNA molecules[5]. Experimentally, RNA structures are determined by techniques like X-ray crystallography, Nuclear magnetic resonance (NMR), cryo-electron microscopy, and chemical probing. These experimental techniques are expensive and have protracted experiment periods with low throughput [6]. To overcome this limitation, computational approaches are used as an alternative to predict the secondary structure of the RNA [7], [8].

Accurate computational prediction of RNA secondary structure is a long-standing problem in bioinformatics. To address this problem, many methods have been proposed, which are either based on (i) comparative sequence analysis [9] and evolutionary profile-based [3], [10]; or (ii) single sequence-based prediction [8], [11], [12]. The operational concept of the models in the first category closely resembles homology modeling, wherein the sequences exhibit a high degree of similarity. The conserved regions of base pairs among homologous sequences are identified through the utilization of pairwise and multiple sequence alignment techniques. The optimal outcomes are achieved when a sufficient amount of homologous sequence data is available and properly aligned.

Some notable works like Pfold [13], RNAalifold [14], PETfold [15], CentroidAlifold [16], and ConsAlifold [9] utilized aligned RNA sequences for their predictions, whereas SPOT-RNA2 [10] and GCNfold [3] used single sequences along with the evolutionary and structural information as features. Since the pool of available RNA sequences remains limited and new RNA families continue to be discovered, relying solely on alignment strategies may not yield satisfactory prediction accuracy. The second category, which is widely employed, involves predicting RNA secondary structures by folding a single sequence using a scoring function as a basis. In the second category, existing works like Mfold [17], CONTRAfold [18], RNAstructure [19], RNAfold [20], and mxfold [21] are based on this minimum free energy (MFE) concept, efficiently implemented through dynamic programming. To reduce the time complexity of dynamic programming-based methods, Linearfold [22] and LinearPartition [23] have proposed an approximate MFE structure in linear time. For the short-range RNAs, the MFE state is applicable. However, the long-range RNAs do not have a stable MFE folding state; rather, they will be stable at a suboptimal energy state. Due to these limitations, this list of diverse methods reaches a plateau and cannot significantly improve the prediction accuracy after reaching a ceiling.

People also tried to improve the accuracy of molecular structure prediction with the use of emerging learning techniques like machine learning and deep learning methods. Some notable works in RNA structure prediction have already used existing deep-learning models to achieve better accuracy [7][24]. The learning-based models can overcome the shortcomings of thermodynamic-based models, as learning-based models do not require prior assumptions, consider both canonical and noncanonical interactions, can handle tertiary interactions and pseudoknots, and can predict previously unrecognized base pairs. Existing methods like SPOT-RNA [12], CDPfold [11], E2Efold [25], Ufold [8], and GCNfold[3] use different learning models like transfer learning, fully convolutional neural networks, and graph convolutional networks to efficiently train the structure prediction model. Though their model shows good prediction accuracy for within-family datasets, but shows very poor performance against cross-family datasets. Only very few RNA sequences have already been discovered, and the majority are yet to be discovered. The prediction model should show fair performance against unseen data. Another shortcoming of the existing models is that they only consider RNA sequences up to a certain length; in most cases, it is up to 600 nucleotides. For longer RNA, they simply omitted or cut to any particular length and then fit into the model. Cutting RNA sequences at arbitrary lengths can lead to the loss of important structural information, as RNA molecules can vary greatly in length, often spanning thousands of nucleotides. Long-range interactions in such sequences frequently include pseudoknots, a special type of non-nested base pairing that contributes to RNA structural stability and occurs predominantly in long RNA molecules (Further details about pseudoknots are given in our supplementary file section S1). However, most existing methods either restrict sequence length or overlook pseudoknots altogether, despite their critical role in maintaining the overall architecture of many RNA families. Further,due to the limited availability of data for RNA compared to proteins, existing large deep learning architectures may encounter challenges such as overfitting and vanishing gradient problems, leading to unsatisfactory outcomes.

To overcome these limitations, we propose a more generalized and efficient deep learning model, rpcFold, for predicting RNA secondary structure from a given RNA sequence. Initially, the RNA sequence is transformed into a two-dimensional (2D) image-like representation with 16 channels, each representing one of the 16 possible base-pairing types, including both canonical and non-canonical interactions, as well as long-range dependencies. Additionally, we incorporate base-pairing possibilities between nucleotides using a Gaussian-based weighting function based on localized pairing rules. To enhance learning efficiency and representation capacity, rpcFold integrates residual connections with parallel convolutional blocks, enabling simultaneous extraction of diverse structural features. Given that structural complexity increases with RNA length, we explicitly evaluate short- and long-range sequences separately. A sliding-window processing scheme is employed to efficiently handle long RNAs, substantially reducing computational overhead and eliminating excessive padding.

Our model is evaluated across both within-family and cross-family RNA datasets, with a separate analysis on long RNAs dominated by pseudoknots—a structural feature often omitted in existing approaches. Compared to both traditional energy minimization approaches and recent learning-based methods, rpcFold achieves 1.73% higher F1 scores on within-family datasets and 4.27% higher F1 scores on cross-family datasets for short-range sequences, relative to the best existing model (UFold). For long-range RNA sequences containing complex pseudoknots, rpcFold attains a 71.1% F1 score in joint prediction of nested and non-nested interactions, a range often overlooked by existing methods. Furthermore, all inter- and intra-dataset redundancies are removed to ensure genuine generalization and avoid inflated performance. Overall, rpcFold provides a straightforward and robust solution capable of accurately predicting canonical and non-canonical interactions, including complex pseudoknots across a wide range of RNA lengths, demonstrating strong generalization capability and enhanced robustness.

## 2. Materials and methods

### 2.1 Dataset preparation

In our study, we used RNA from three well-known benchmark datasets: RNAstralign [26], ArchiveII [27], and the bpRNA_1m dataset [28]. In the RNAstralign and ArchiveII datasets, RNA sequences are well-separated based on their RNA families. The details of each of these datasets are given in Supplementary Tables S1 and S2. The bpRNA_1m contains 102318 RNA sequences from 2588 different families, ranging between 1nt and 4381nt. The length-wise distribution of RNA sequences across each dataset is provided in Supplementary Figures S3 and S4, while the RNA family-wise length distribution histograms for each dataset are separately presented in Supplementary Figures S5 and S6.

For any kind of deep learning-based model, the data must be unambiguous and non-redundant. Thus, for our dataset, we remove all the incomplete sequences and any nucleotide except standard adenine(A) / uracil(U) / guanine(G) / cytosine(C). The RNA fragments contribute incomplete knowledge of the structures and mislead the model training. Therefore, we have excluded sequences that do not form any structure (remain single-stranded), or if they are only part of stems or loops, sections of an RNA structure, or peptide chains, particularly those sequences shorter than 30 nucleotides. Also, we removed all the duplicate and similar sequences from each dataset. To have an impartial comparison with the existing methods, which contain some degree of duplication, only the cross-dataset duplications in the ArchiveII dataset are retained, while we removed their counterparts from the other datasets. The RNAStralign dataset is widely used as a benchmark in existing studies and contains more accurate structures than the bpRNA_1m dataset. Therefore, keeping the samples with higher accuracy for training the model could enhance its performance. Further, we noticed that bpRNA_1m contains 92% data within the range of 200nt, and the data is highly similar. So, before applying the aforementioned preprocessing steps, we reduced the homology of the bpRNA_1m dataset using the CDHIT-EST with a threshold of 0.9. The steps of data preprocessing are described in detail in Figure 1.

**Figure 1.**
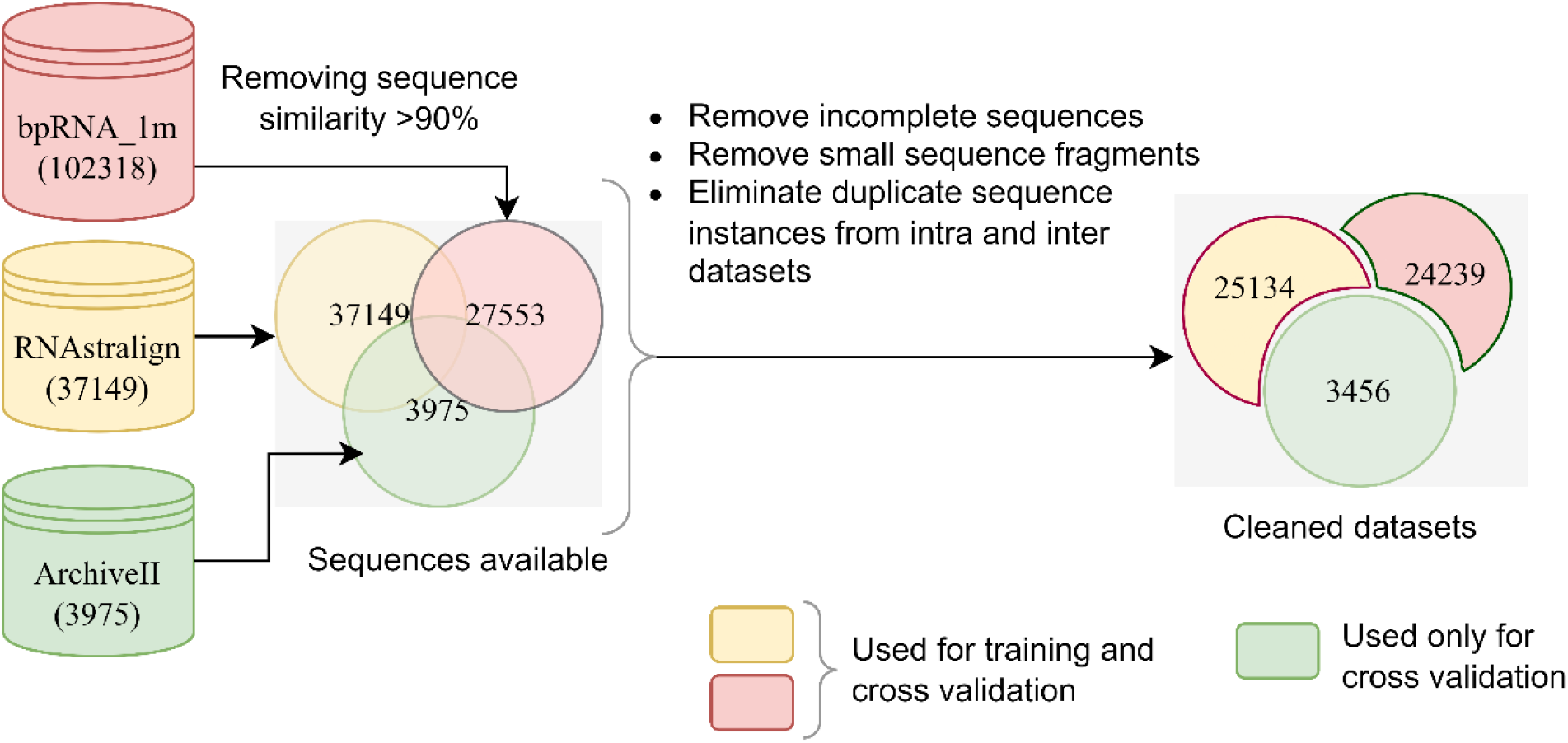
The data processing flow, where the number of RNAs after each step is labelled and color-coded for easy visualization.

For fair performance comparison of our model, we divide our processed RNAstralign and bpRNA_1m datasets into a training set and test set, using family-wise stratified sampling with a ratio of 4:1. Further, to confirm the robustness of our model against unseen data, we test our model using the ArchiveII dataset, which has never been used for training. We employ a stratified sampling approach to ensure that RNA sequences from all families, as well as subfamilies in the dataset, are proportionally represented in both the training and test sets. A brief overview of available families and subfamilies for the RNAstralign dataset is given in Table 1.

**Table 1.** RNA family-wise details of the pre-processed RNAstralign dataset.

| RNA Family | RNA subfamilies | Count | Min sequence length | Max sequence length |
| --- | --- | --- | --- | --- |
| 16S RNA | Actinobacteria, Alphaproteobacteria, Aquificae, Bacilli, Mollicutes | 9550 | 54 | 1829 |
| 5S RNA | Archaea, Bacteria, Eukaryota, Mitochondria, Plastids | 8558 | 106 | 132 |
| group_I_intron_database | IA1, IA2, IA3, IB1, IB2, IB4, IC1, IC3, IE, IE1, IE2, IE3 | 679 | 163 | 615 |
| tRNA_database | - | 6347 | 64 | 95 |

### 2.2 Problem Statement / Input and Output Representation

Given an RNA sequence of length *N* as input, *S* = (*s*_1_, *s*_2_, *s*_3_ … *s*_*N*_) where *s*_*i*_ ∈{‘A’, ‘U’, ‘G’, ‘C’}. We have to predict the RNA secondary structure output that is in the form of *T* ∈ {0,1}^*N*×*N*^ a binary matrix (as represented in Figure 2) where,

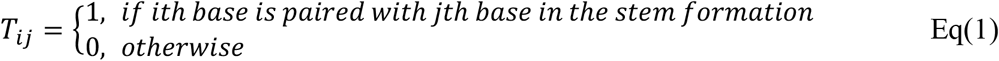

**Figure 2.**
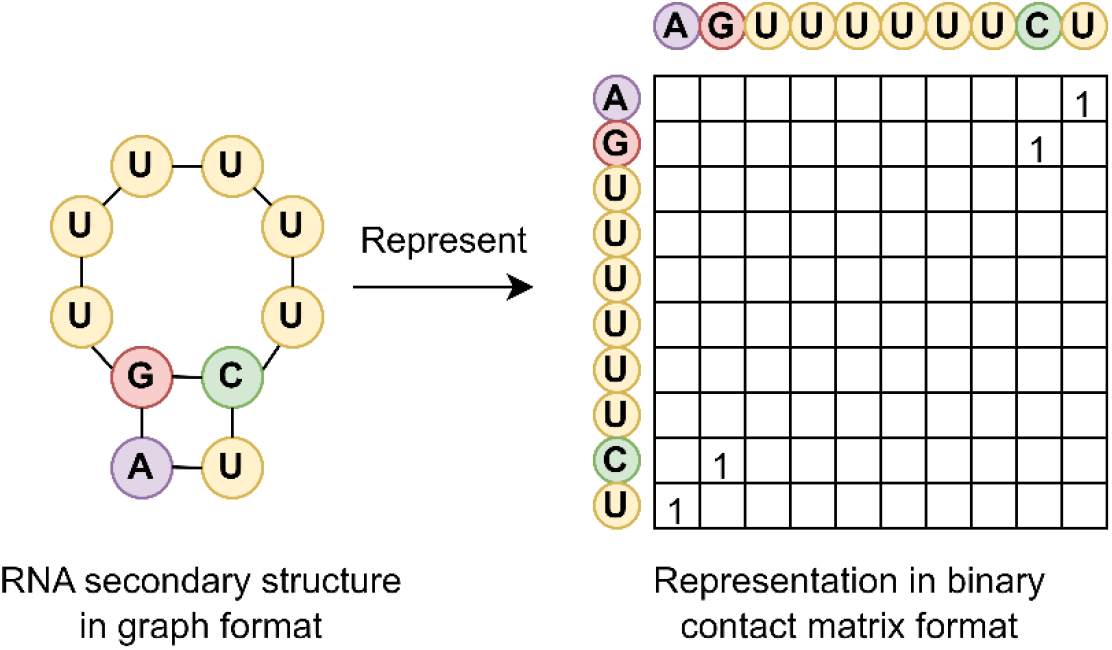
Mapping RNA secondary structure graphs to matrix representations. Empty cells are assigned a value of 0.

Traditional deep learning approaches often represent RNA as a sequential input and utilize RNN-based models, such as LSTMs and GRUs, to learn long-range base interactions. Recently, transformer architectures, convolutional models such as U-Net, and hybrid combinations of both, which support parallel computation and more effectively capture global dependencies, have also been used. However, despite these advancements, such models are often constrained by several drawbacks, like they typically require large numbers of trainable parameters, are computationally intensive, and are data-hungry, making them less suitable for tasks involving limited and imbalanced RNA datasets. Moreover, models relying on sequential architectures still face efficiency bottlenecks during training and inference, especially when processing long RNA sequences. Our model is built on a parallel convolutional neural network architecture enhanced with residual blocks, enabling effective learning of base-pairing interactions. The integration of residual connections helps mitigate issues such as vanishing gradients and supports deeper network training. Additionally, a sliding window mechanism is introduced for handling long-range RNA sequences, which reduces computational complexity and facilitates efficient processing of long RNA inputs without sequence truncation or excessive padding.

In our approach, short-range and long-range RNA sequences are handled separately. Short-range sequences are fed directly into the model, while long-range sequences are divided into overlapping subsequences using a sliding window mechanism. Specifically, a sequence of length *n* is split into ((*n* − *w*)/*d*) + 1 subsequences, where *w* is the window size, and *d* is the length of the slide. This process, illustrated in Step 1 of Figure 3, allows us to process RNA sequences of arbitrary length efficiently, without imposing length restrictions or incurring excessive computational overhead.

**Figure 3.**
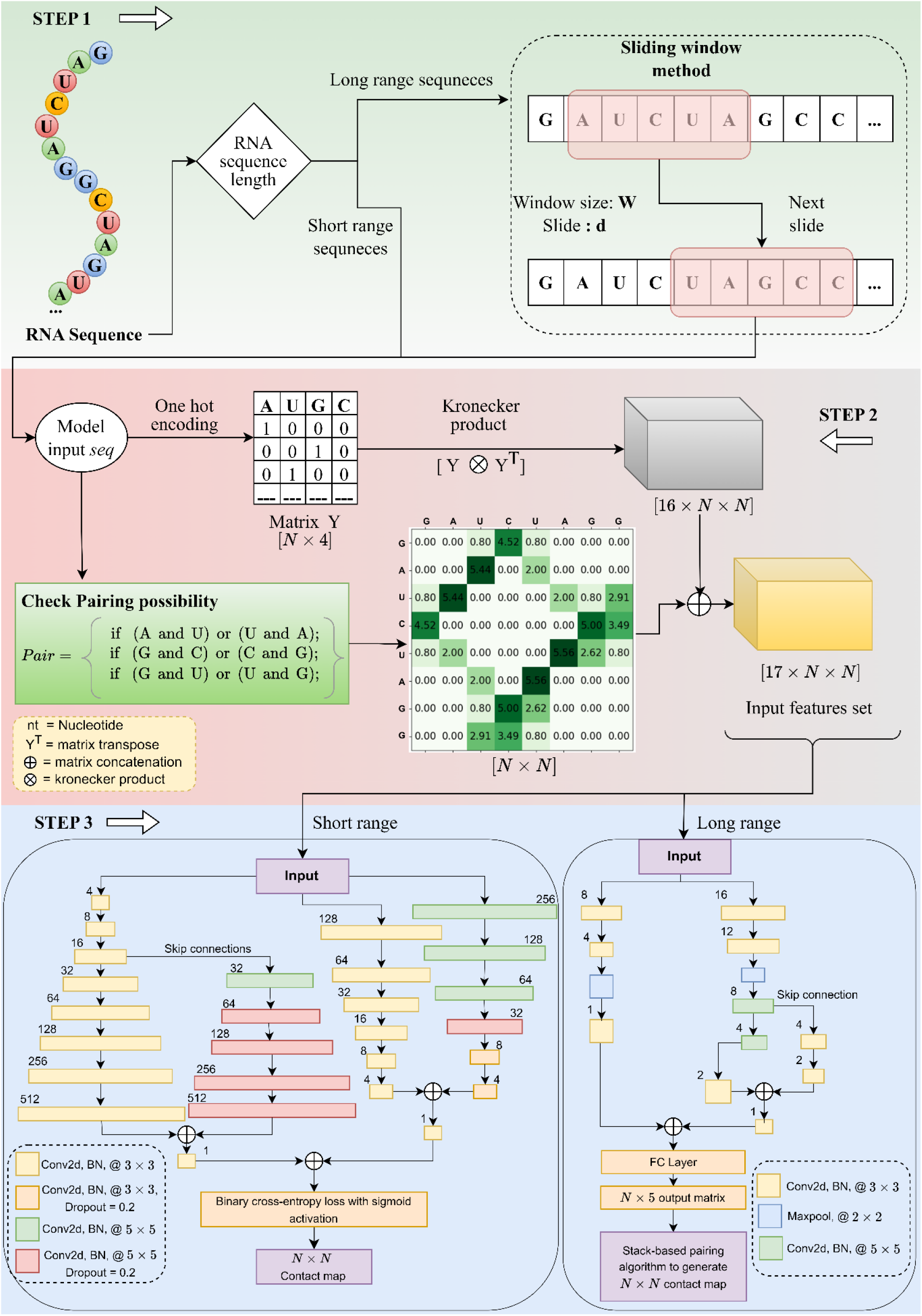
The overall architecture of our model. Step 1(green background): The input sequence is separated based on the length. For each long RNA sequence, sub-sequences will be generated using the sliding window method, and each of the sub-sequences is then passed to step 2 for calculating the feature set. The small sequences are directly passed to step 2 without subsequence generation. Step 2 (pink background): Here, we calculate the pairing possibility matrix based on three given rules. We also calculate all 16 possible base pairs using the Kronecker product and utilize them as features. After concatenating both features, we have 17 × *N* × *N* an input feature set that is passed to our model in Step 3. Step 3 (blue background): For small and long-range sequences, we proposed two different approaches. Model 1 is designed for smaller sequences, consisting of parallel convolutional layers and producing output directly in the *N* × *N* format. Model 2 is for longer sequences, which takes data in a window-wise manner and transforms the learned features into the *N* × 5 output format using the fully connected layer. From that, we achieve our desired *N* × *N* output using a stack-based pairing algorithm inspired by the pushdown automata (PDA).

In step 2, we convert each of our sequences (the subsequences of long RNAs) into a *Y* ∈ {0,1}^*N*×4^ binary matrix using one-hot encoding representation (details are provided in the Supporting Information S2).

#### Algorithm 1

The algorithm to compute the base pairing possibility matrix W for a given input RNA sequence

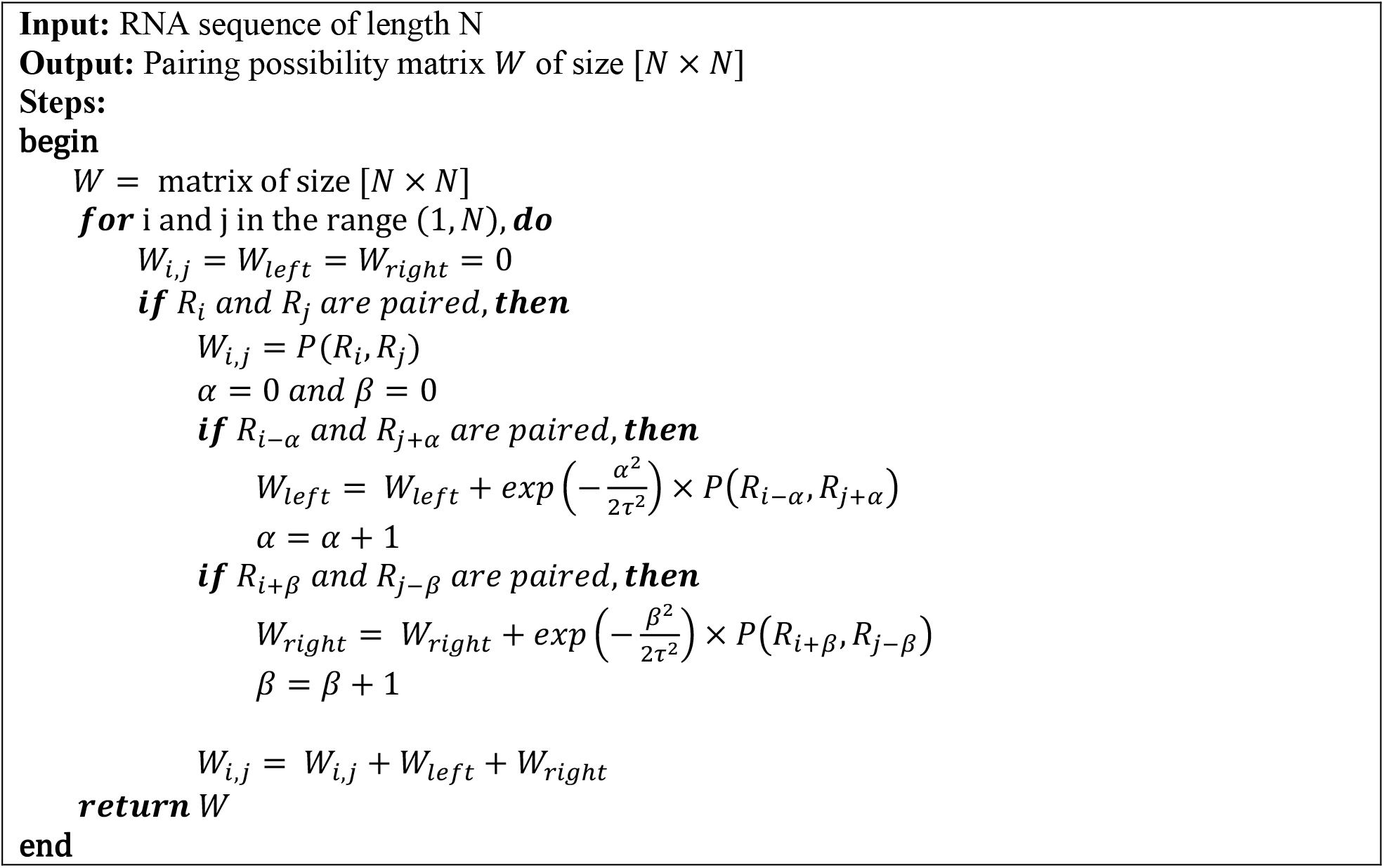

Next, to deal with the short-range as well as long-range base interactions, our data is transformed into an image-like representation using the Kronecker product. The Kronecker product on the matrix *Y* and its transpose *Y*^*T*^ gives us a [4*N* × 4*N*] matrix representation. To consider all the possible base pairs, the matrix is reshaped into a [16 × *N* × *N*] matrix, which is treated as an image of resolution [*N* × *N*] image with 16 channels.

To reduce the sparsity of our proposed matrix, we introduce the second feature, where we consider the pairing possibilities between each base separately. Here, instead of one-hot encoding, we used the concept from CDPFold [11] to encode the input sequence. The input RNA sequence of length N is converted into a matrix *W* ∈ [*N* × *N*], based on the algorithm as discussed in Algorithm 1.

At first, the matrix *W* is initialized to zeros. Then for each base *R*_*i*_ and *R*_*j*_, we calculate the base pairing possibility between them. Here *P*(*R*_*i*_, *R*_*j*_) denotes the pairing weight based on the number of hydrogen bonds between those two bases. The calculation of pairing weights for two bases *R*_*i*_ *and R*_*j*_ at any two positions *i and j*, is shown in Eq2.

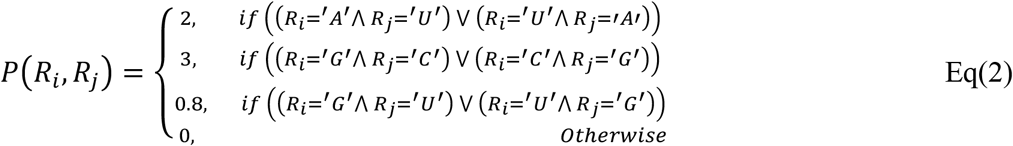

For any two positions *i* and *j* in the RNA sequence, we consider whether the corresponding bases form a pair within a stem region, as such pairings contribute significantly to the structural stability. To capture this, we evaluate the pairing status of neighboring bases surrounding *i* and *j*, since central base pairs within a stem tend to exhibit greater stability than those at the ends. To incorporate this biological insight into our model, we employ a Gaussian weighting function to assign higher importance to base pairs located near the center of stems, effectively modeling local stability variations. This results in a position-specific pairing possibility matrix *W* of size [1 × *N* × *N*], which, when combined with the Kronecker product, results in the final input feature vector of size [17 × *N* × *N*].

Here, the image-like input representation offers several advantages. First, it helps the explicit modeling of long-range interactions by transforming the RNA sequence into a two-dimensional format, allowing local base-pairing patterns between distant nucleotide segments to be effectively captured. Second, it does not differentiate between canonical and non-canonical pairing; rather, it uniformly considers all possible base-pairing combinations. Moreover, since base-pairing information is directly extracted from input.ct files, the representation also incorporates non-nested structural elements such as pseudoknots, enhancing the structural completeness of the input.

### 2.3 Deep learning network architecture

To accommodate RNA sequences of varying lengths, we propose two deep learning architectures, as illustrated in Step 3 of Figure 3. Both models consist of a parallel two-dimensional convolutional neural network (2D-CNN) architecture along with skip connections to capture all the intricate canonical and non-canonical base pairing details of the RNA structures.

Our first model deals with short-range interactions. It consists of parallel convolutional pathways. Each pathway processes the input tensor using a series of convolutional layers with varying numbers of filters, kernel sizes, and configurations to extract multiscale features. The input format of this model is in [*N* × *N* × 17] format. The upper pathway employs a progressively increasing number of filters (4 →8 → … → 512), while the lower pathway uses a decreasing filter configuration (256 → 128 → … → 4). This design allows the network to extract hierarchical features at both coarse and fine resolutions. Each convolutional layer is followed by a ReLU activation function and batch normalization layers. Some convolutional blocks also incorporate dropout (rate = 0.2) to enhance generalization. To achieve network resilience and stabilize the training by preventing the exploding gradient problem, skip connections are introduced, which allow earlier feature representations to be merged with deeper layers, promoting richer feature fusion. After the parallel branches process the input, their outputs are merged using element-wise addition operations. The final feature map is passed through a sequence of Conv2D layers with decreasing filter sizes (4 → 1), and produces the final model output, which is a logit-valued contact matrix *T* ∈ {0,1}^*N* ×*N*^. To handle the strong class imbalance in this contact matrix prediction, where positive base pairs are much fewer than negative (not paired) ones, we add a weighted binary cross-entropy loss, applied directly to the raw output logits. Specifically, the BCEWithLogitsLoss is used, which combines a sigmoid activation internally with the binary cross-entropy formulation as shown is eq 3 and 4, ensuring numerical stability during optimization.

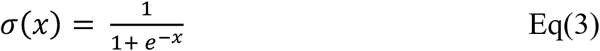

Where x is the unbounded output from the previous layer. A positive class weight of *w* = 300 is assigned to the positive class to amplify the penalty for misclassified pairs and guide the model toward higher recall of true positives. So our final loss function is defined as-

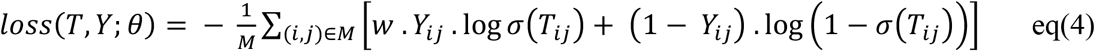

Where θ represents all parameters in the neural network, w is the weight added to the positive class, *M* represents the set of all valid base pairs within the actual sequence length *N, Y* ∈ (0,1)^*N*×*N*^ denotes the ground truth, and *σ*(*T*_*ij*_) is the sigmoid-activated prediction for an N base pair (*i, j*).

For handling long-range interactions, we employ a parallel 2D-CNN architecture, followed by a max-pooling layer and a fully connected layer. The input to this model is formatted as a tensor of shape [*W* × *W* × 17] format where *W* is the window size. After testing multiple window sizes, we found *W* = 200 to yield the best performance. The network processes overlapping windows across the full sequence, and the outputs are concatenated to produce an *N* × 5 matrix, where *N* is the sequence length. This matrix is decoded into an *N* × *N* contact map using a stack-based parsing algorithm inspired by pushdown automata (PDA).

Our model can derive short-range interactions very efficiently and provide highly accurate and robust prediction results. After incorporating the sliding window concept along with our model, the ensemble model shows better performance for long RNA structure prediction as it can capture the long-range interactions very accurately. Now, our model can handle RNA of any length. This window-based formulation significantly reduces the effective computational cost and memory usage, as the model no longer operates over the full *N* × *N* space but instead focuses on smaller *W* × *W* regions where *W* ≪ *N*.

### 2.4 Post-processing

After obtaining the symmetric contact scoring matrix from our model, we apply a constraint-guided post-processing procedure to extract the final RNA secondary structure. We integrate biological domain knowledge to ensure that the predicted contact map adheres to biologically feasible. To this end, we enforce the following four hard constraints:

i. Symmetry: The contact matrix must be symmetric.
ii. Base Pair Validity: Only canonical (A-U, G-C) and wobble (G-U) base pairs are permitted, although this constraint can be relaxed depending on the dataset characteristics.
iii. Loop Length: No sharp loops are allowed by enforcing *Y*_*ij*_ = 0, ∀*i, j such that* |*i* − *j*| < 4, ensuring a minimum loop length of 3.
iv. Non-overlapping Pairs: Each nucleotide can pair with at most one other base, i.e., 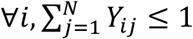

To implement constraints (ii) and (iii), we define a binary mask matrix *M*(*s*) as follows in eq 5:

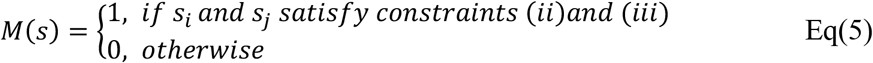

We then refine the model output *T* by enforcing symmetry and element-wise masking using the Hadamard product (denoted by °) as shown in eq 6:

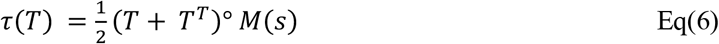

Here *T*^*T*^ denotes the transpose of matrix T. This formulation ensures that τ(*T*) is symmetric and compliant with constraints (i)–(iii).

To address the last and most important constraints, i.e. not allowing any overlapping pairs, we relax the problem into a linear programming optimization as eq 7:

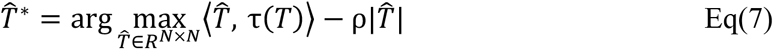

Here, the goal is to find an optimal scoring matrix 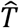 that maximizes similarity to τ(*T*) (measured via the Frobenius inner product) while enforcing sparsity through the *L*_1_ norm penalty term. The hyperparameter *ρ* controls the sparsity level, effectively regulating the number of predicted base pairs. Finally, the binary contact map 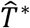 is obtained by thresholding 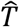 using an offset value, which is selected through grid search.

## 3. Results

### 3.1 Overview of rpcFold

We develop a model capable of handling RNA sequences of varying lengths. Our analysis indicates that the majority of RNA sequences in the bpRNA_1m and RNAstralign datasets fall within the short-range category, while long-range sequences are relatively less frequent. These long-range sequences often contain complex non-nested base pairs, such as pseudoknots, which are particularly difficult to predict. To address this, we divide the dataset based on sequence length and train separate models for short-range and long-range RNAs, enabling more effective learning of the distinct base-pairing patterns present in each category. Further performance evaluation and detailed analysis for both canonical and non-canonical structures (primarily pseudoknots) are provided in this Results section.

To benchmark rpcFold, we conduct three experiments: (1) training on the RNAstralign training set and testing on its test set, (2) evaluating on the ArchiveII dataset without retraining, using the model trained on RNAstralign, and (3) training and testing on the bpRNA_1m dataset.

### 3.2 Performance evaluation

RNA secondary structure prediction is a classification problem[29], where a base can be unpaired or paired. Therefore, we evaluated the performance of our RNA secondary structure prediction model through precision, recall, F1-score, and accuracy, which are defined as

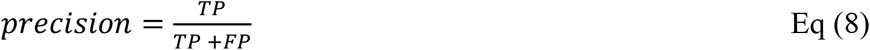

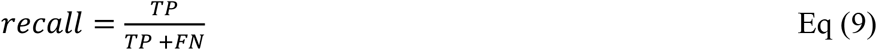

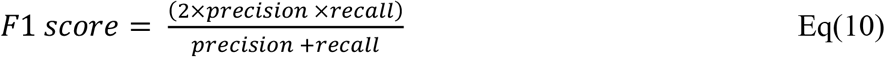

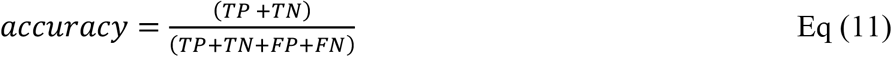

Base pairs that are both predicted by the model and present in the reference (accepted) structure are considered true positives (TP). In contrast, predicted pairs that do not appear in the accepted structure are categorized as false positives (FP). Similarly, base pairs that are present in the accepted structure but not predicted are considered false negatives (FN). Finally, base pairs that are neither predicted nor present in the accepted structure are classified as true negatives (TN). F1-score is the harmonic mean of precision and recall, and can handle the uneven class distribution. Here, the number of true negatives is extremely high due to the quadratic growth of possible base pairs with sequence length. Since actual base pairs are sparse, this inflates accuracy and makes it misleading. Therefore, we evaluate our model performance using precision, recall, and F1 score metrics.

### 3.3 Experimental results on RNAstralign and ArchiveII dataset

In this section, we report results on the RNAstralign and ArchiveII test sets.

We partition the RNAstralign benchmark dataset into training and testing subsets using family-wise stratified sampling in a 4:1 ratio, ensuring proportional representation of RNA sequences from all families. For long-range interaction experiments, only the Group I intron and 16S rRNA families are considered, while short-range experiments include all available families. The short-range model is evaluated on sequences of length ≤600 nucleotides, aligning with typical constraints (≤500–600 nt) used in most existing methods. For both short- and long-range experiments, we perform 5-fold cross-validation using datasets containing 16,666 and 8,462 samples, respectively.

To ensure a fair comparison, we retrain all learning-based baselines (UFold, MXfold2[30], and E2Efold) on our training data and evaluate them on our test set. For traditional energy-based approaches, predictions are generated using their official software or publicly available web servers. As shown in Table 2, state-of-the-art methods achieve F1 scores ranging from 0.42 to 0.965, whereas our model attains an F1 score of 0.981. Learning-based models often exhibit lower recall than precision, while energy-based models tend to favor recall; in contrast, our approach maintains a balanced trade-off, achieving high and closely matched precision and recall values.

**Table 2.**
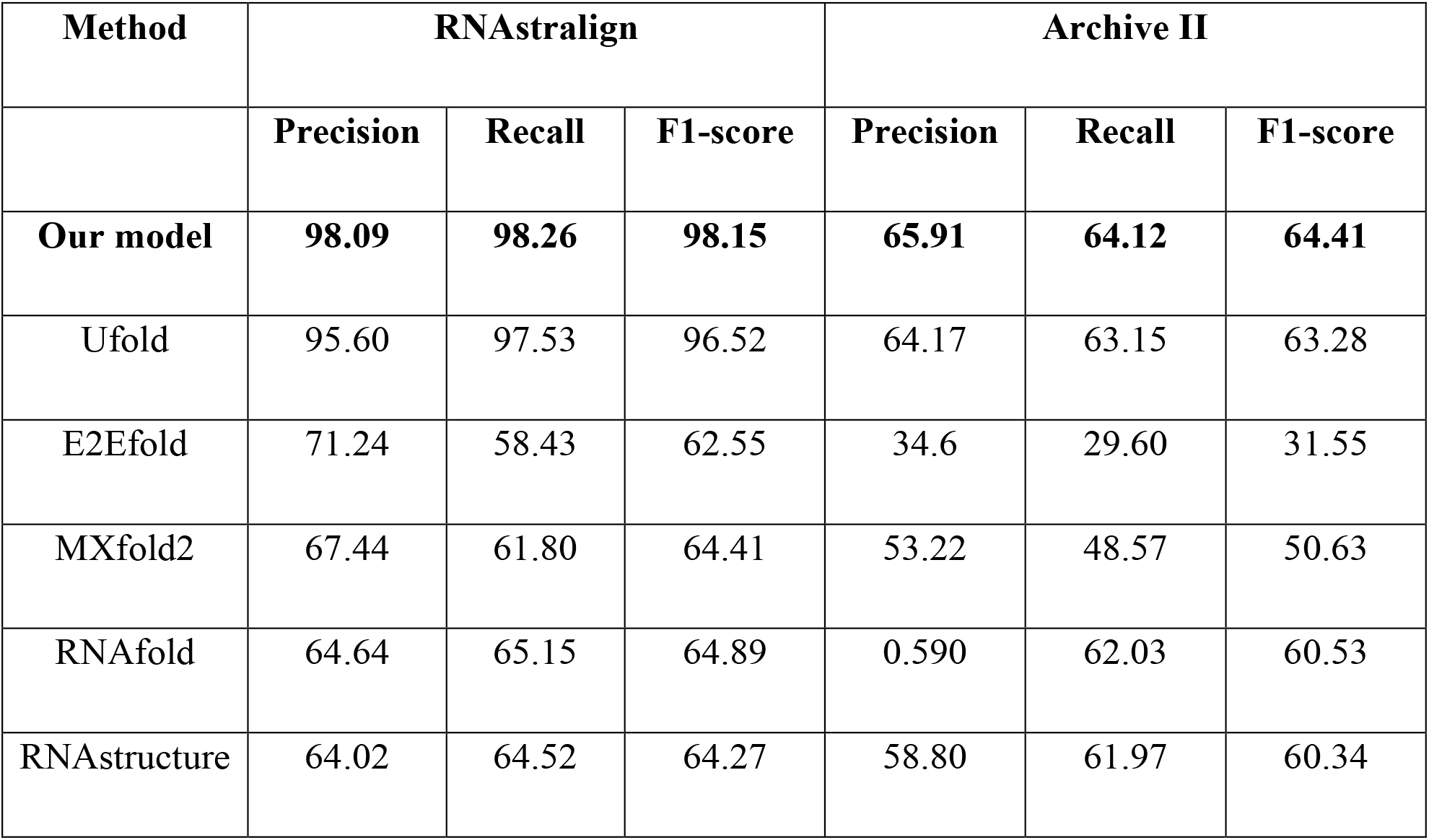
Performance evaluation and cross-validation on our RNAstralign and Archive II test datasets. Bold values indicate the best performance achieved for each metric in (%).

We further assess generalization capability by evaluating on the ArchiveII dataset as an unseen test set. The model, trained solely on the RNAstralign training set, is applied directly to ArchiveII without retraining. To prevent performance inflation, we remove all redundant and highly similar sequences between the two datasets, keeping them only in ArchiveII and excluding them from RNAstralign (as described in Section Dataset Preparation). This ensures no sequence-level leakage between training and evaluation. Unlike many existing learning-based approaches that overlook inter-dataset redundancy, our setup provides a more realistic estimate of generalization.

All learning-based baselines are retrained on our non-redundant RNAstralign training set and evaluated on the full ArchiveII dataset, without excluding any RNA family—even if absent in the training set. Overall results are summarized in Table 2, while Supplementary Table S3 reports family-wise performance, including whether each family was present during training. Unlike prior studies that restrict evaluation to seen families, we include both seen and unseen families in our assessment. Although this leads to lower absolute F1 scores than previously reported, it offers a fairer and more robust comparison. Certain RNA families—such as RNase P, SRP, tmRNA, and telomerase—are absent from the training set and show low F1 scores across all models. However, rpcFold consistently achieves higher scores on these unseen families than competing methods, highlighting its robustness and capacity to generalize to novel RNA families.

### 3.4 Experiments on the bpRNA_1m dataset

Next, the bpRNA-1m dataset is divided into training and test subsets, and the corresponding experimental results are presented in Table 3. Compared to existing state-of-the-art (SOTA) methods, our model achieves superior performance. As with earlier evaluations, we retrain the learning-based baseline models using our bpRNA-1m training set and evaluate them on the corresponding test set to ensure a fair comparison.

**Table 3.**
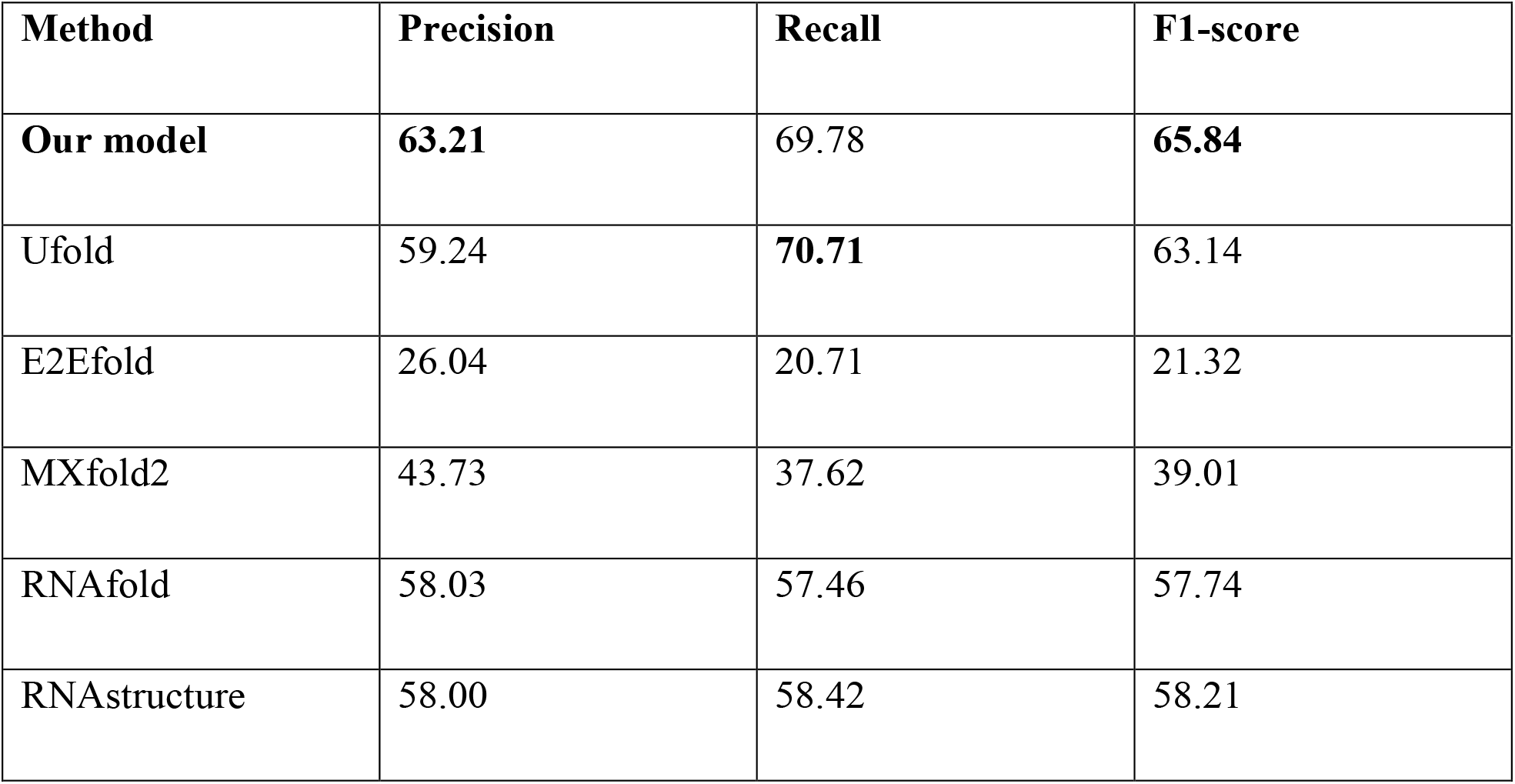
Prediction on bpRNA_1m test set for cross-validation. **Bold** values indicate the best performance achieved for each metric (in %).

Notably, our model exhibits a minimal gap between precision and recall, significantly smaller than that observed in other SOTA approaches. This indicates that the model handles both false positives and false negatives effectively.

It is also important to highlight that bpRNA-1m comprises RNA sequences from over 2,000 different families, making structure prediction considerably more challenging compared to the previous two datasets. Achieving an F1 score of 0.658 under these conditions is promising for a learning-based model, especially one that relies solely on sequence input without incorporating prior structural assumptions or thermodynamic features.

### 3.5 Experimental results on Long-range interactions

Further, we evaluate our long-range interaction prediction model for predicting base-pairing information in longer RNA sequences (length >600 nt), most of which contain pseudoknots. As suggested by Mathews [29], RNA structure prediction can be assessed not only by exact base pair positions, but also by considering ±1 neighborhood positions. Given the complexity of longer sequences, we combine and report the prediction performance of exact and ±1 base pair positions.

Table 4 lists the goodness of our prediction when tested on only pseudoknots, only canonical pairs, and combined. Since jointly predicting pseudoknots and canonical pairs in a single sequence remains a highly challenging task, the observed performance (F1 score ≈ 0.711 indicates that our model demonstrates promising capability in handling such complex structures. We report performance only on the RNAstralign dataset, since the other datasets lack sufficiently long sequences. We do not compare our model with existing works in this setting, as learning-based models typically restrict RNA lengths to 500∼600 nucleotides, while methods like RNAfold and RNAstructure do not account for pseudoknots.

**Table 4.** Prediction on RNAstralign test set for sequences with pseudoknot (>600 length). (in %)

|  | Only pseudoknots | Pseudo+canonical pairs | Only canonical pairs |
| --- | --- | --- | --- |
| **Precision** | 66.82 | 71.62 | 80.45 |
| **Recall** | 56.92 | 70.57 | 82.52 |
| **F1-score** | 61.47 | 71.16 | 81.47 |

## 4. Discussion

The starting approaches in the RNA structure prediction domain are mainly based on thermodynamics-based properties. Afterward, researchers incorporated thermodynamic features along with learning techniques for better learning of their model. However, most thermodynamics-based models rely on prior assumptions—such as favoring minimum free energy (MFE) structures, considering only canonical base pairs (A–U, G–C, and sometimes G–U), excluding pseudoknots, and using predefined thermodynamic parameters derived from experimental data. These models typically evaluate the stability of individual stems and loops to estimate the overall free energy and infer the most stable structure. In contrast, we avoid such assumptions entirely and instead rely solely on RNA sequence features, enabling a more general and flexible model that is not restricted by thermodynamic constraints and can be applied broadly across diverse RNA families and structural complexities.

During our analysis, we observed that prior evaluations on commonly used benchmark datasets may have been influenced by inter-dataset redundancy, potentially leading to inflated performance estimates. In particular, the ArchiveII dataset shares 1,893 exact sequence matches with the RNAstrAlign dataset, introducing bias into the reported performance metrics when used for cross-validation. To ensure a more reliable and fair assessment, we carefully removed all redundant sequences across datasets. Specifically, we excluded exact sequence duplicates from the RNAstralign training set while retaining them only in the ArchiveII test set. Upon retraining several state-of-the-art (SOTA) models on this cleaned, non-redundant dataset and evaluating them on the curated ArchiveII test set, we observed a noticeable performance drop across all models. This drop is largely due to the absence of overlapping RNA families between training and test sets, resulting in models encountering truly unseen families during evaluation. Some earlier approaches attempted to address inter-dataset redundancy; however, when evaluating on the ArchiveII dataset, they included only those RNA families that were present in the training set, omitting unseen families. This selective evaluation introduces a performance bias, as it favors familiar RNA families and does not reflect the model’s ability to generalize to novel or unseen families. Further, some existing models worked with only the RNAstralign dataset, which comprises sequences from only eight RNA families, and were evaluated on the closely related and redundant ArchiveII dataset. While this setup can yield high training and test accuracy, it may not reflect true generalization. When such models are applied to more diverse datasets like bpRNA_1m—which includes sequences from over 2,400 RNA families—their performance often drops significantly. This highlights the importance of developing models that can generalize well across both narrow and broad family distributions. To ensure robust and unbiased evaluation, we eliminate both inter- and intra-dataset redundancies, strictly separate training and test sets, and report performance across within- and cross-family scenarios to demonstrate the generalizability of our model.

Here, we proposed our model, which can handle both short-range and long-range RNA sequences using the advantage of both skip connections along with parallel convolutional layers. The skip connection helps in more efficient and stable learning by vanishing the gradient problem during training, whereas the parallel convolutional layers have the potential for capturing diverse features simultaneously, leading to better performance on sequence feature extraction tasks. We also introduce the sliding window method, which gives us the flexibility to handle RNA sequences of any length without illogically breaking them in between or without adding unnecessary padding, which increases the computational cost. The sliding concept also helps the model capture local patterns and dependencies and facilitates the extraction of relevant features from input sequences. Our model can predict both the nested and non-nested pseudoknot base pairings present in RNA molecules. The performance prediction on inter-family and cross-family datasets reveals that our model shows good generalizability. Compared to existing methods, rpcFold improves F1 scores by 1.73% and 4.27% on the within-family and diverse bpRNA_1m datasets over the best-performing model, and by 51.23% and 13.05% compared to the second-best model.

Additionally, we compared the prediction accuracy of our rpcFold model with existing tools, MXfold2 and RNAfold, using the RNA sequence with ID DQ170668.ct (Bacilli, 16sRNA) from the RNAstralign test set. This RNA sequence, consisting of 824 nucleotides, includes 216 nested base pairs and 15 non-nested pseudoknots. Figure 4 illustrates the ground truth secondary structure alongside the predictions made by rpcFold, RNAfold, and MXfold2. Notably, our rpcFold model demonstrates a strong capability to predict pseudoknots accurately. It achieves an overall F1 score of 0.823 for predicting all the base pairs, and an F1 score of 0.896 specifically for pseudoknot prediction, highlighting its effectiveness in handling complex RNA structures. RNAfold and MXfold2 cannot predict pseudoknot structures.

**Figure 4.**
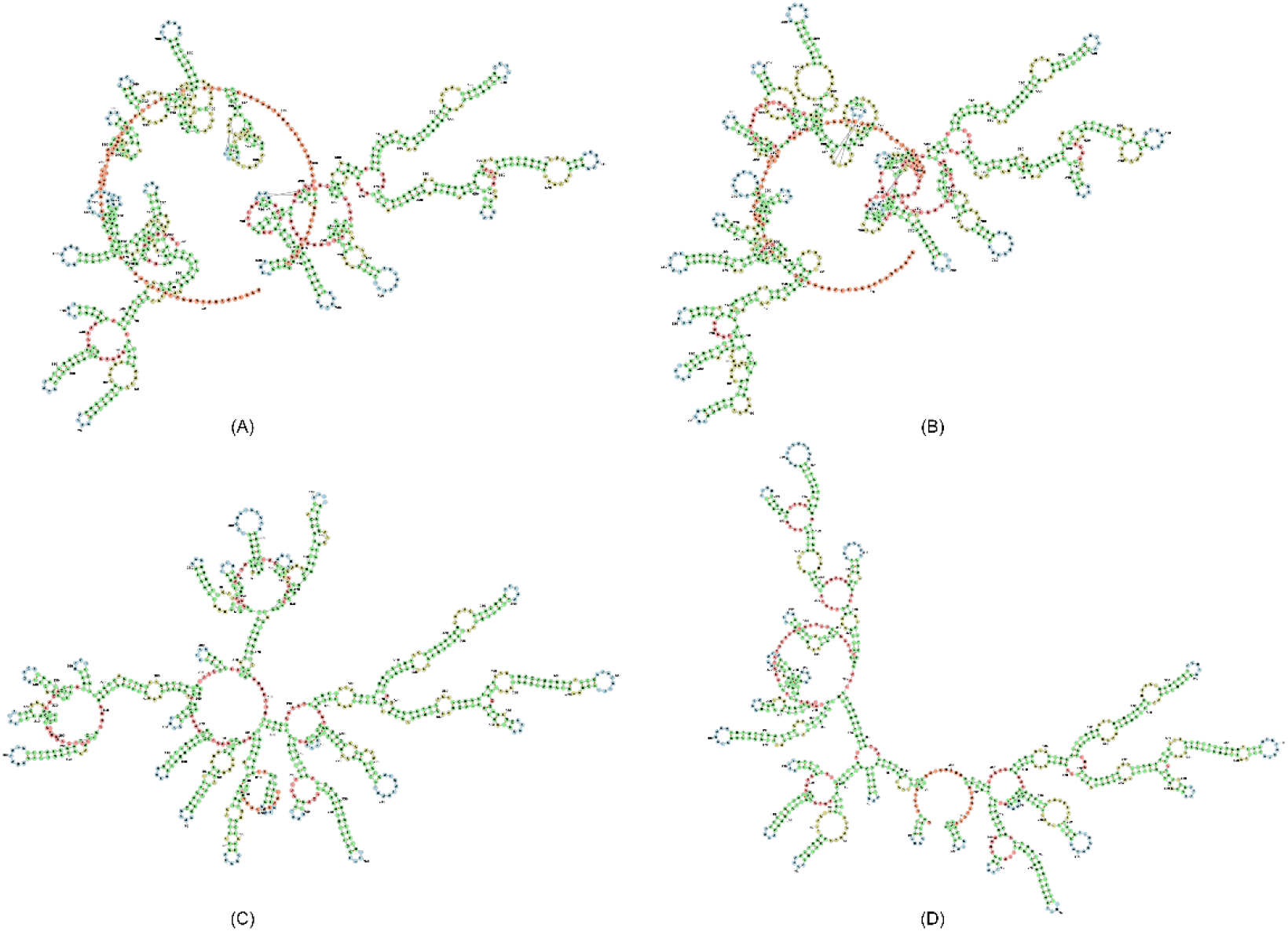
Prediction comparison of rpcFold, RNAfold, and MXFold2 with native structure on RNA ID: DQ170668.ct, (Bacilli, 16sRNA) from RNAstralign database where sequence length 824nt. (A) Native structure, (B) predicted structure by rpcFold with 0.823 F1-score, (C) predicted structure by RNAfold with 0.671 F1-score, (D) predicted structure by MXFold2 with 0.730 F1-score.

## 5. Conclusion

We propose a novel algorithm for predicting the secondary structure of noncoding RNAs. Given the wide variation in RNA sequence lengths from a few nucleotides to several thousand, we employ a residual parallel convolutional neural network architecture that effectively captures both short-range and long-range base interactions. To address the challenges associated with processing long sequences, we incorporate a sliding window approach, which allows efficient handling of long sequences without breaking or excessive padding, as is commonly seen in existing methods. Our model relies solely on sequence information extracted from established benchmark datasets. The raw RNA sequences are transformed into an image-like representation that captures all potential base pairing combinations. Additionally, we construct a second feature matrix that encodes base-pairing possibilities based on local structural context (e.g., stem and loop regions), using a Gaussian-based weighting function. These two feature matrices are then concatenated and fed into the model for training. In this work, we explicitly consider special non-nested base pairs known as pseudoknots, and report their prediction performance separately to demonstrate our model’s effectiveness in handling such complex structures. When evaluated on standard benchmark datasets, rpcFold demonstrates consistent improvements over state-of-the-art RNA secondary structure prediction methods. Specifically, on the RNAstralign, ArchiveII, and bpRNA_1m test sets, rpcFold surpasses the best existing model by 1.65%, 1.73%, and 4.27% in F1 score, respectively, and outperforms the second-best model by 51.23%, 6.46%, and 13.05%, respectively. Additionally, it exhibits improved performance across diverse RNA families, highlighting its robustness and generalizability. Given the data-driven nature of deep learning models, we recognize that future improvements can be achieved by incorporating high-quality RNA sequences from a broader range of families. Overall, our approach offers valuable contributions to the field of RNA secondary structure prediction, providing a reliable framework that can be leveraged and extended by researchers addressing similar challenges.

## Acknowledgment

We do hereby acknowledge the use of the Supercomputer facility Paramshakti of the Indian Institute of Technology Kharagpur.

## Authors

Nandita Sharma received the B.Tech. degree from the University of Burdwan and the M.Tech. degree from NIT Patna, India. She is currently pursuing a Ph.D. degree in the Centre for Computational and Data Sciences at IIT Kharagpur under the supervision of Dr. Pralay Mitra. Her research focuses on RNA structure and functional analysis.

Pralay Mitra received the B.Sc. (with Hons.) degree in physics, and the B.Tech. degree in computer science and engineering from the University of Calcutta, Kolkata, India, in 1999 and 2002, respectively, and the M.E. degree in computer science and engineering from the Indian Institute of Engineering Science and Technology, Shibpur, Howrah, India, (formerly Bengal Engineering College and Science University, Shibpur), in 2004, and the Ph.D. degree from the Indian Institute of Science, Bangalore, Bengaluru, India, in 2010. Since 2005, he is actively working on Bioinformatics and Computational Biology. He has developed a number of algorithms for protein-protein docking, predicting protein assembly from the crystal structures, and protein design. He was a Research Associate with the Indian Institute of Science, Bangalore from 2010 to 2011 and Research Fellow with the University of Michigan, Ann Arbor, MI, USA from 2011 to 2013. Currently, he is an Associate Professor with the Department of Computer Science and Engineering, Indian Institute of Technology Kharagpur, India.

## Authors Contribution

N.S.: Conceptualization, Data curation, Formal analysis, Investigation, Methodology, Software, Validation, Visualization, Writing – original draft, Writing – review and editing

P.M.: Conceptualization, Formal analysis, Investigation, Methodology, Resources, Supervision, Writing – original draft, Writing – review and editing

