## Supplementary Tables S1, Supplementary Figures S3, Supplementary Figures S5 and S6, Supplementary Table S3 for "rpcFold: Residual Parallel Convolutional neural network to decipher RNA folding from RNA sequence"

### Table of Contents


| Section | Page No |
|---|---|
| S1. Details about the stem and loop regions in an RNA secondary structure | 3 |
| S2. Details of the input features | 4 |
| S3. Supplementary figures | 5 |
| S4. Supplementary tables | 7 |
| S5. References | 10 |

### S1. Details about the stem, nested and non-nested loop regions in an RNA secondary structure

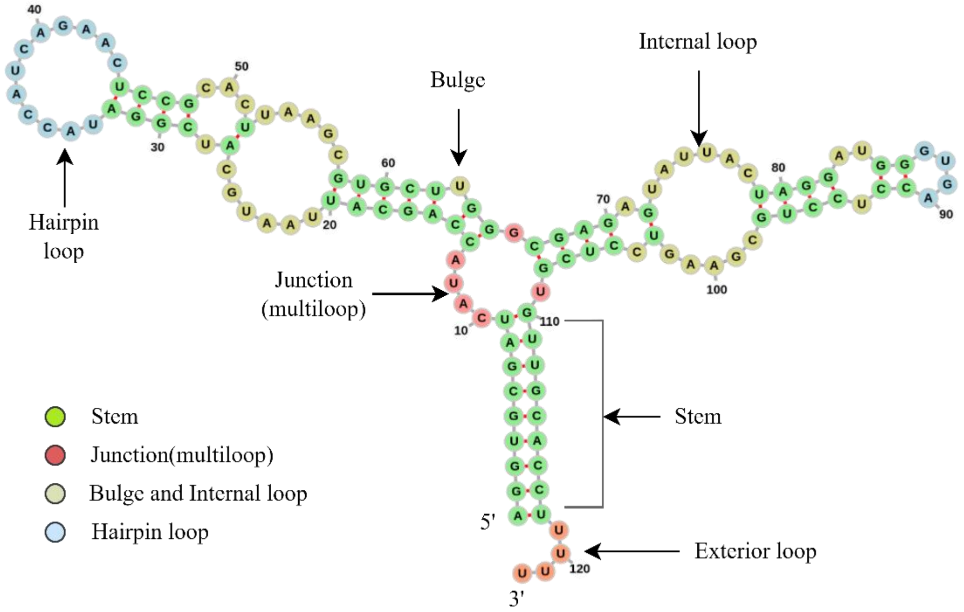


**Sequence:** AGGUGCGAUCAUACCAGCAUUAAUGCAUCGGAUACCAUCAGAACUCCGCACUUAAGCGU GCUUGGGCGAGAGUAUUACUAGGAUGGGUGACCUCCUGCGAAGUCCUCGUGUUGCACCUUUUU

**Structure:** (((((((((....(((((((......(.((((............))))...).....))))).)).((((.(......((((..((....)).))))......).)))).)))))))))....

**Figure S1:** Details about stem and loops in the secondary structure of an RNA, taken from RNAStralign dataset with RNA ID: E02541.ct (Eukaryota, 5sRNA).

RNA secondary structure mainly contains the hydrogen bond information between the base pairs. Here the non-paired bases are denoted as part of the loop and the paired bases form the stem. Based on the position, loops can be of different types. A bulge and internal loop exist in between the paired base(stem) region. Whereas a hairpin loop exists at the end of any stem. A junction (multiloop) is found as a connector between different stems. An exterior loop is a point where if we cut the structure then we get two nonoverlapping substructures. The idea about these substructures is shown in Figure S1, will help us to analyse the efficacy of the predicted RNA structure. One of the ways to represent the secondary structure of the RNA is the dot-bracket notation where stem regions are indicated as paired regions using opening '(' and closing ')' brackets. Whereas the loop regions are dented with dot '.' values. The sequence and the corresponding secondary structure in dot-bracket notation of the RNA ID: E02541.ct from RNAStralign dataset is also shown in the Figure S1.

A pseudoknot is a type of RNA non-nested loop structure that arises when bases in a loop pair with complementary bases outside the loop, leading to crossing interactions between base pairs as shown in below Figure S2. Unlike standard nested base pairings, pseudoknots involve non-nested configurations—formally, when two base pairs $(i, j)$ and $(k, l)$ satisfy the condition $i < k < j < l$. These structures contribute significantly to the structural and functional complexity of RNA molecules. Pseudoknots are known to play critical roles in various biological processes, including ribosomal frameshifting, the activity of ribozymes, and viral genome replication. Due to their complex topology, they pose challenges for computational RNA structure prediction, as many traditional algorithms are designed to handle only nested base pairs. Despite this, accurate modeling of pseudoknots is essential for understanding RNA function and improving the reliability of structure-based predictions in RNA biology.

We handle the pseudoknots and non-pseudoknot sequences separately in our work.

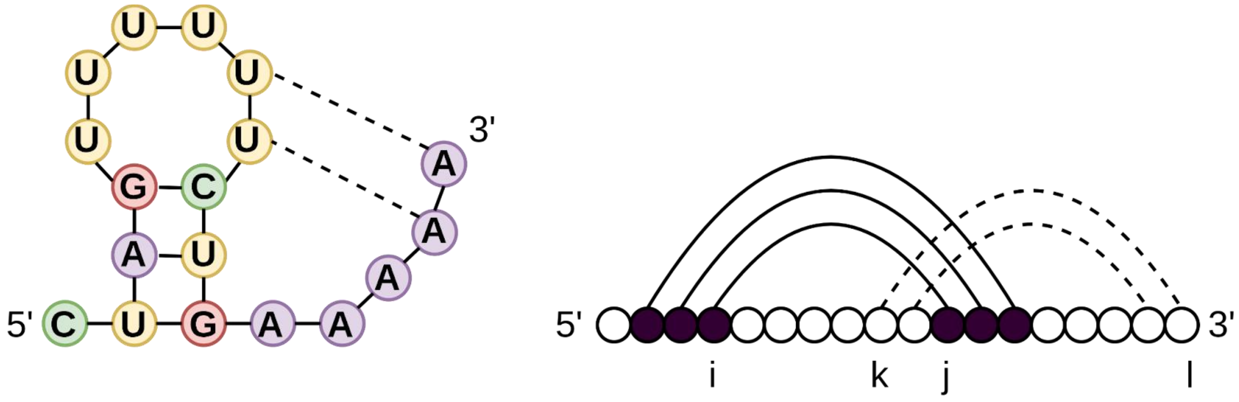


.

**Figure S2:** Graphical representation of pseudoknots in RNA secondary structure.

#### S2. Details of the input features

In our rpcFold model, we have passed two features extracted from input sequence directly. Suppose, $S = (s_1, s_2, s_3 \dots s_N)$, where $N$ is the length of the input RNA sequence and $s_i$ ∈{'A', 'U', 'G', 'C'}. For preparing the first feature, we encode each of the bases into a vector form using one-hot encoding as shown in Eq S1.

$$Y_i = \begin{cases} [1,0,0,0], & where\ s_i='A' \\ [0,1,0,0], & where\ s_i='U' \\ [0,0,1,0], & where\ s_i='G' \\ [0,0,0,1], & where\ s_i='C' \end{cases} \qquad \text{(Eq S1)}$$

After getting the above binary matrix, we perform Kronecker product on $Y$ and $Y^T$ (Y transform) and get the results in a $[4N \times 4N]$ matrix representation. To consider all 16 possible base pairs in RNA, we convert our input into a $[16 \times N \times N]$ matrix which is similar to an image representation of a $[N \times N]$ image with 16 channels.

To reduce the sparsity of our proposed matrix and to add features like pairing possibilities between bases, the concept CDPFold[1] is used here. Since each of the nucleotides in RNA is affected by its neighboring nucleotides, we calculate the base pairing possibility for each nucleotide in a sequence. A pair base in the middle of a stem is relatively more stable than a pair base at the end of a stem. To consider this structural information, the concept of local weighted linear regression is used. We calculate the non-binary matrix $[1 \times N \times N]$ which can potentially reduce the model's sparsity and offer additional base pairing information.

After concatenating both the feature matrices we finally get our $[17 \times N \times N]$ input feature matrix and pass it to our deep learning rpcFold model for model training.

### S3. Supplementary Figures

We have RNA sequences of varying lengths in the bpRNA_1m[3], ArchiveII[4], and RNAStralign[2] datasets. A violin plot depicting the length wise distribution of these RNA sequences is shown in Figure S3.

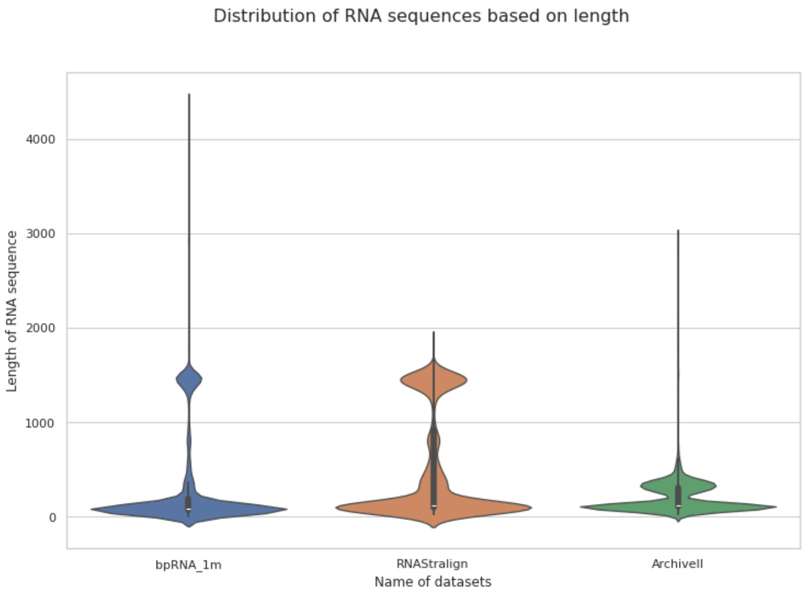


**Figure S3:** Visualization of the distribution of RNA sequences on different datasets length-wise.

We also present a histogram plot and distribution of the RNA sequences based on length for each dataset in Figure S3. Figures S3 and S4 demonstrate that the majority of the sequences are smaller in size, comprising nearly 80% of the dataset.

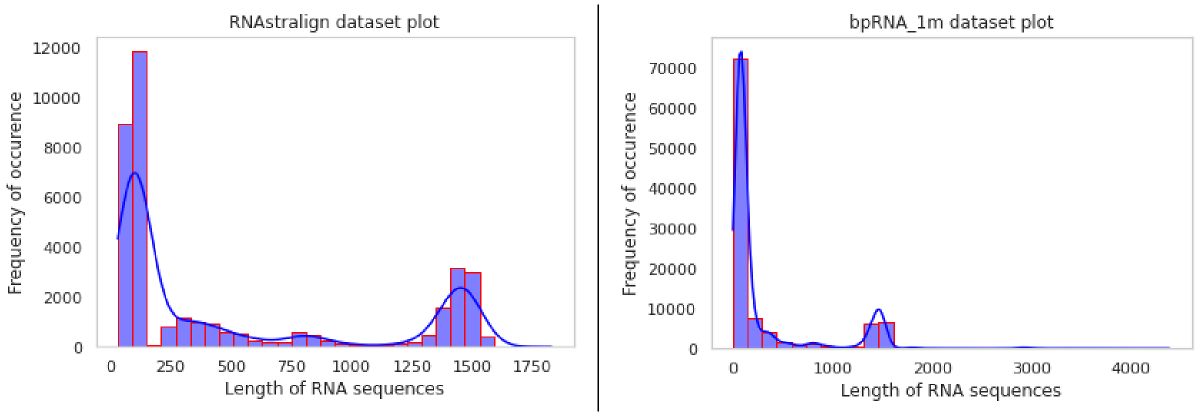

| (A) | (B) |
| --- | --- |

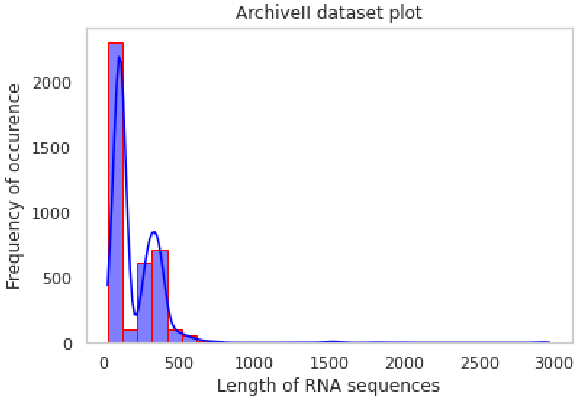


(C)

**Figure S4:** Distribution of RNA based on length on (A) RNAStralign[2], (B) bpRNA_1m[3], and (C) ArchiveII[4] dataset.

Further, the length-based histogram plot and distribution of the RNA sequences family-wise is shown separately for different datasets in supplementary Figures S4 and S5 below.

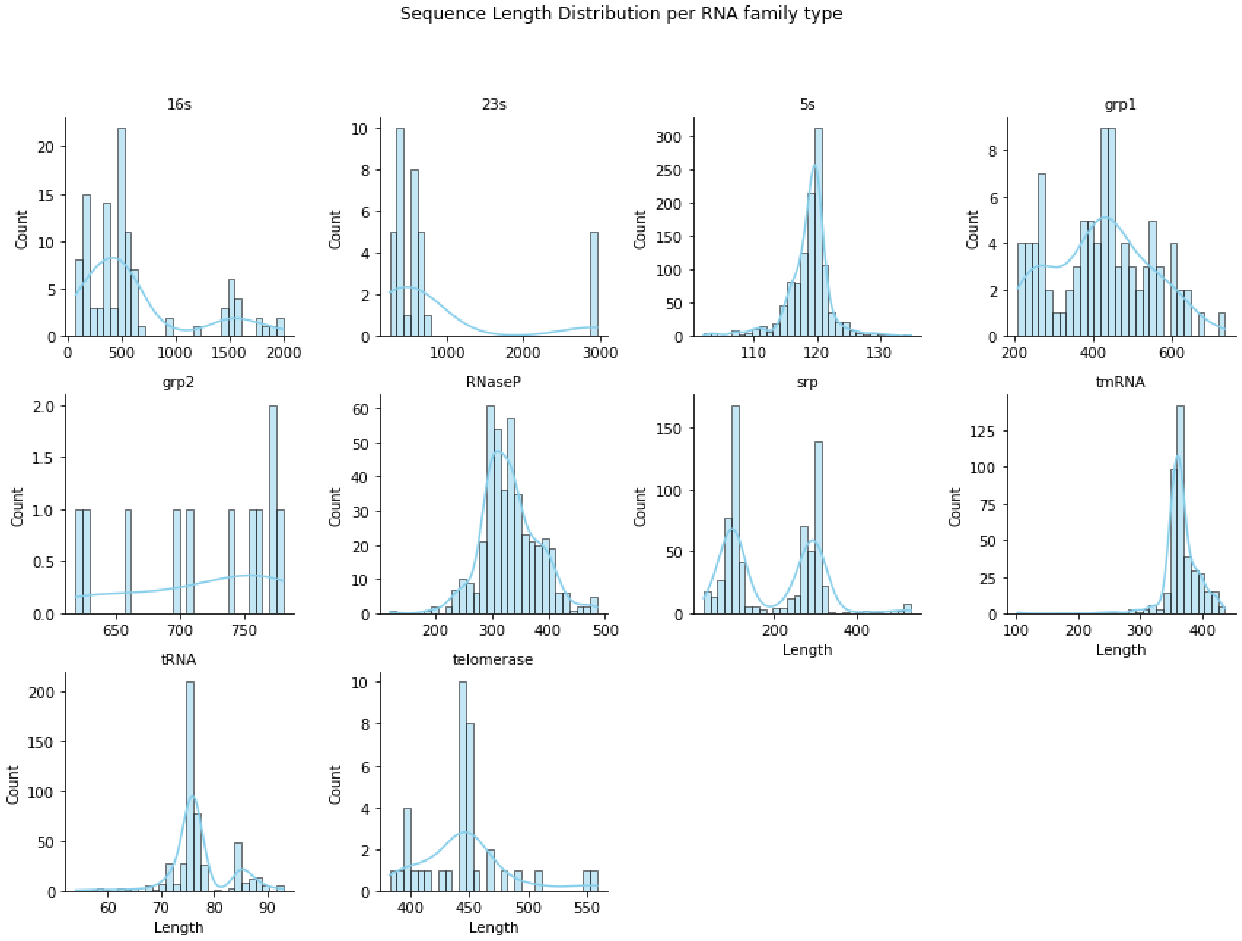

**Figure S5:** Length-wise distribution of RNA sequences across different RNA families in the ArchiveII dataset.

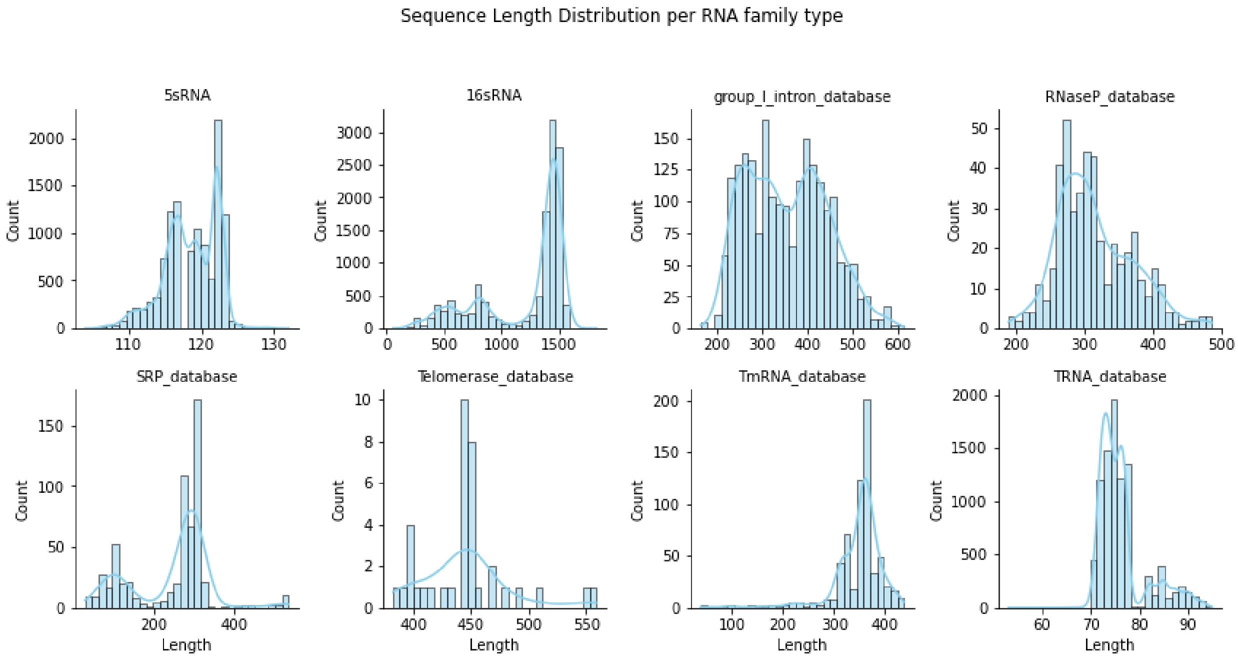


**Figure S6:** Length-wise distribution of RNA sequences across different RNA families in the RNAstralign dataset.

### S4. Supplementary Tables

The details of RNAstralign and ArchiveII datasets, including their Family wise count, RNA length ranges are given in below Supplementary table S1 and S2.

**Table S1:** The RNAstralign dataset statistics

| RNA Family | Count | Min sequence length | Max sequence length |
|---|---|---|---|
| 16sRNA | 12608 | 54 | 1829 |
| 5sRNA | 11419 | 104 | 132 |
| RNasep | 467 | 189 | 486 |

| | | | |
|---|---|---|---|
| SRP | 601 | 30 | 533 |
| tRNA | 9245 | 53 | 95 |
| tmRNA | 637 | 39 | 437 |
| telomerase | 37 | 382 | 559 |
| group-I intron | 2135 | 163 | 615 |

**Table S2:** The ArchiveII dataset statistics

| RNA Family | Count | Min sequence length | Max sequence length |
|---|---|---|---|
| 23sRNA | 35 | 242 | 2968 |
| 16sRNA | 109 | 73 | 1995 |
| 5sRNA | 1147 | 102 | 135 |
| RNAsep | 429 | 120 | 486 |
| SRP | 699 | 28 | 533 |
| tRNA | 492 | 54 | 93 |
| tmRNA | 404 | 102 | 437 |
| telomerase | 37 | 382 | 559 |

| | | | |
|---|---|---|---|
| group-I intron | 93 | 210 | 736 |
| group-I intron | 11 | 619 | 780 |

**Table S3:** Performance evaluation of the ArchiveII dataset family-wise.

| Functional- class | No. of RNA samples | F1 score | | | Family present on Train set |
|---|---|---|---|---|---|
| | | Our Model | UFold | E2Efold | |
| 16sRNA | 80 | 0.570 | 0.590 | 0.136 | Yes |
| 23sRNA | 21 | 0.334 | 0.332 | 0.018 | No |
| 5sRNA | 1147 | 0.974 | 0.974 | 0.843 | Yes |
| RNasep | 429 | 0.459 | 0.431 | 0.046 | No |
| group1_intron | 83 | 0.613 | 0.641 | 0.027 | Yes |
| srp | 699 | 0.231 | 0.190 | 0.025 | No |
| tRNA | 492 | 0.963 | 0.966 | 0.089 | Yes |
| tmRNA | 404 | 0.312 | 0.306 | 0.019 | No |
| telomerase | 35 | 0.168 | 0.152 | 0.014 | No |
